# Recurrently Mutated Genes Differ between Leptomeningeal and Solid Lung Cancer Brain Metastases

**DOI:** 10.1101/222349

**Authors:** Yingmei Li, Boxiang Liu, Ian David Connollyt, Bina Wasunga Kakusa, Wenying Pan, Seema Nagpal, Stephen B. Montgomery, Melanie Hayden Gephart

## Abstract

**Purpose:** Brain metastases from non-small cell lung cancer (NSCLC) engraft and grow either within the brain (solid) or diffusely on its surface (leptomeningeal disease; LMD). Routine clinical diagnostics have low sensitivity and provide no information about the underlying mutations. A recurrent mutation analysis of LMD and a comparison between solid and LMD NSCLC brain metastases have yet to be explored.

**Experimental Design:** We performed whole-exome sequencing (WES) on eight cerebrospinal fluid (CSF) specimens from NSCLC LMD patients. We compared our LMD sequencing data with a published data set of 26 NSCLC solid brain metastases to determine the relative mutation frequency. We then performed a retrospective chart review of an additional set of 44 NSCLC LMD patients to further evaluate LMD mutations and clinical prognosis.

**Results:** Six (75%) LMD cases had mutations in *EGFR*, while none had *KRAS* mutations. Retrospective chart review revealed only 4 LMD cases (7.7%) with *KRAS* mutations, but 33 cases (63.5%) with *EGFR* mutations. *TP53* was mutated in 4/8 LMD (50%) cases and 13/26 of solid metastasis (50%). The median interval for developing LMD from NSCLC was shorter in EGFR-mutant (16.3 mo) than wild-type (23.9 mo) patients (p = 0.017).

**Conclusions:** *EGFR* and *TP53* mutations were frequent in LMD exomes (combined frequency 87.5%), suggesting that PCR-based mutation detection assays towards these two genes could be a useful complement to current diagnostics. Correlations of *EGFR* in LMD and *KRAS* in solid metastases suggest molecular distinctions or systemic treatment pressure underpinning differences in growth patterns within the brain.

**Translational Relevance:** Leptomeningeal disease is a diffuse, malignant, and incurable metastatic brain tumor that accounts for 5-10% of brain metastases. Patients with LMD do not undergo biopsy and their overall prognosis is poor (median survival 3 to 27 months), making it difficult to collect sufficient samples for recurrent mutation analysis. Standard diagnostic procedures (MRI and cytology) for LMD provide no genetic information. To understand the mutation landscape of LMD, we performed whole-exome sequencing on eight lung-derived LMD cases. We showed that mutations in *EGFR* occurred more frequently in LMD than solid brain metastases, but *KRAS* mutations were not present in LMD. Further, mutations in recurrent genes such as *EGFR* and *TP53* could be reliably detected in CSF via droplet digital PCR. Targeted analysis of recurrent mutations thus presents a useful complement to the existing diagnostic toolkit, and differences in mutations between LMD and solid brain metastases suggest distinct molecular mechanisms for growth.

## Introduction

Brain metastases most commonly arise from lung, breast, and skin cancer. The development of brain metastases denotes a very poor prognosis, with a median survival from 3 to 27 months.(1) The incidence of brain metastases is highest in lung cancer, with 10-25% of patients found to have brain metastases at the time of diagnosis, and 40-50% of lung cancer patients developing brain metastases later during the course of their disease.(2) There are two distinct subtypes of brain metastases, solid metastasis to the brain parenchyma, and leptomeningeal metastasis (LMD) to the covering of the brain and cranial nerves. Although solid lung-to-brain metastases frequently respond well to surgery and radiation, LMD does not have a durable treatment.(3–5) Many clinical trials specifically excluded NSCLC patients with LMD or any history of brain metastases, making it challenging to ascertain the effects of treatment specifically on brain metastases.(6) Treatment options for LMD include radiation, chemotherapy and targeted therapy, yet these are generally ineffective, with a median survival of less than 6 months.(7,8) Due to its diffuse nature, surgical resection or biopsy is not an option in LMD. The diagnosis is made by MRI or via lumbar puncture, which may capture a few cells present in cerebrospinal fluid (CSF). The rarity of cases, rapid disease progression, and lack of available tissue for research has severely limited our understanding of LMD genetics and biology. The genetic underpinnings that differentiate LMD from solid tumor metastases are not clear; to our knowledge, a cohort analysis on recurrently mutated genes in lung cancer LMD has yet to be reported. Prior studies showing the genomic evolution of primary cancers and solid brain metastases(9–11) have uncovered some genes that appear to be important in brain metastases.(12–15) Despite this, it is unknown whether specific mutations in lung cancer enhance its ability to thrive in the leptomeninges or within the brain. We hypothesized that LMD harbored a distinct mutation profile compared to solid brain metastases, allowing these two metastatic subtypes to thrive in different brain niches. To this end, we compared our whole-exome profiles of a NSCLC LMD cohort to previously sequenced, publicly available solid lung-to-brain tumor metastases, in order to discover differences in their mutational landscapes.

We obtained tumor DNA from cytology-positive CSF samples and DNA from matched normal tissues to perform exome sequencing and variant calling. Figure 1 depicts the overall workflow of our study. A total of 5 CSF samples with matched normal samples (blood or saliva) and 3 CSF samples without matched normal samples from LMD patients were processed (Table S1). In this study of lung-to-brain metastases, we investigated (i) the similarity/heterogeneity among LMD patients; (ii) genomic distinctions between LMD and solid brain tumors by comparing our data with available data of solid brain tumors;^10^ (iii) detectability of mutations in the cell-free DNA component of CSF; and (iv) the correlation of recurrent LMD mutations with disease progression and survival.

**Figure 1.**
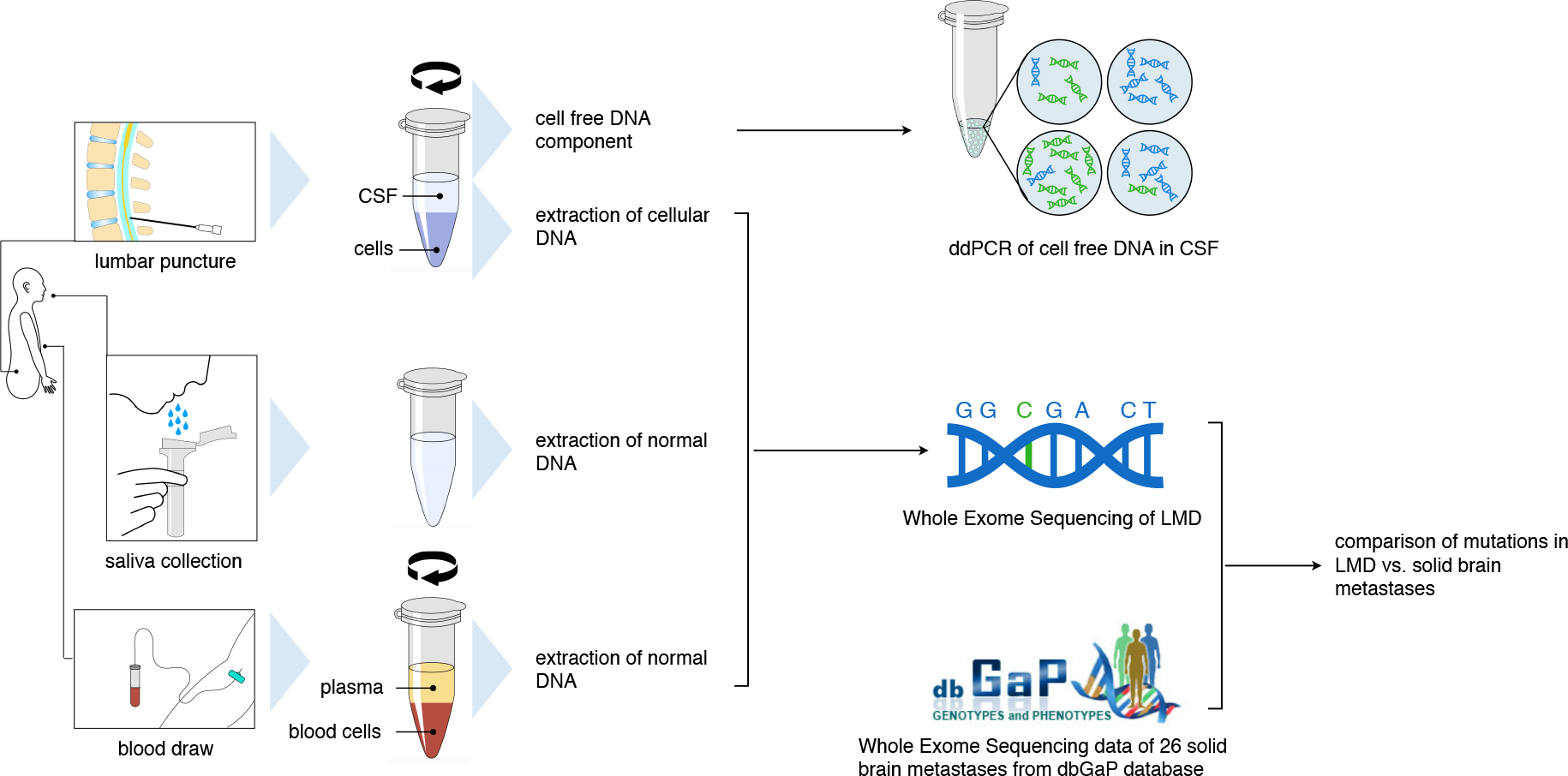
Workflow for comparing solid versus leptomeningeal NSCLC brain metastases. Patients undergoing clinical confirmation of leptomeningeal disease (n=8) have a lumbar puncture compared to normal DNA from either a corresponding blood or saliva sample. Following centrifugation and extraction of DNA, exome sequencing data were compared to dbGaP data on 26 solid NSCLC brain metastases.

## Materials and methods

### Sample Processing

All samples were collected through an Institutional Review Board-approved informed consent process. Only specimens not required for clinical pathological diagnosis were utilized in this study. No procedures were performed for the exclusive purpose of research. CSF and saliva samples were obtained at the time of lumbar puncture in the outpatient clinic. The portion of the CSF that was taken for clinical diagnosis was reviewed in parallel by cytopathology. Additional sequencing data from solid lung cancer brain metastases were obtained from the database of Genotypes and Phenotypes (accession number phs000730. v1.p1).

Blood and CSF were centrifuged (1000 g, 10 min) within 1 h of collection. The initial supernatant from centrifuged blood samples underwent an additional centrifugation step (10,000 g, 10 min) to further separate plasma and blood cells. The supernatant and pellet components were transferred to separate tubes and stored at -80 degrees C until ready for DNA extraction. Saliva was collected through the use of a commercially available kit (Oragene). Tumor tissue was collected and snap frozen at -80 degrees C within an hour of removal. DNA from CSF was extracted with a QIAamp DNA Mini Kit (Qiagen), while DNA from blood cells and tumor tissue was extracted with a DNEasy Blood and Tissue Kit (Qiagen). Saliva DNA was extracted using a reagent prepIT L2P (genotek). Extracted DNA was fragmented to approximately 300 base pairs by sonication (S220 focused ultrasonicator, Covaris).

### Whole Exome Sequencing

Indexed Illumina libraries were created (KAPA Hyper Prep Kit, Kapa Biosystems) from 20 ~ 100 ng of CSF cell pellets DNA, blood cellular DNA and saliva DNA. Exonic DNA was captured using NimbleGen SeqCap EZ Human Exome Library (Roche). Approximately 100 million reads per library were sequenced with 150-bp paired end runs on a NextSeq sequencing system (Illumina).

### Solid Brain Metastasis Data

We downloaded publicly available data collected by Brastianos *et al*.^10^ We used sam-dump to convert SRA files to alignment files for 26 solid NSCLC brain metastases in the dataset.

### Variant Calling

Prior to alignment, we trimmed Illumina adaptors using trimmomatic v0.36 with the following parameters: “PE -threads 8 -phred33 TRAILING:5 SLIDINGWINDOW:4:20 MINLEN:35 CROP:100.” We aligned each pair of FASTQ files using bwa mem v0.7.12 with default parameters. We used Picard v1.111 to sort, index, and mark duplicate in the SAM files. We used GATK to perform indel-realignment, base quality score recalibration with default parameters. We called germline variants using GATK HaplotypeCaller with “-stand_call_conf 30 -stand_emit_conf 10” and restricting genomic interval to those in the Roche SeqCap EZ Exome v3 capture targets. We filtered SNP using GATK’s VariantFiltration with "QD < 2.0 ║ FS > 60.0 ║ MQ < 40.0 ║ MQRankSum < -12.5 ║ ReadPosRankSum < -8.0".

### Somatic Variant Identification

We identified somatic variants with MuTect2 with dbSNP v138 and COSMIC v54 (20120711) as annotated germline and somatic variants. Calling somatic mutations without normal controls poses a challenge as the somatic mutation call set is confounded with low frequency and germline mutations. For the three samples without normal controls, we adopted a pipeline developed by Hiltemann *et* al.(16) In brief, we compared CSF samples against 433 normal individuals from the 1000 Genomes project,(17) and removed any variants that presented in any normal individuals. Further, we used annovar (version dated 2016-02-01)(18) to remove variants in 1000g2015aug_all, snp138NonFlagged, esp6500siv2_all, and exac03 databases. In order to retain likely somatic mutations, we kept variants found in COSMIC 70 and ICGC. We filtered variants against five normal samples processed using the same library prep and bioinformatics pipeline to remove bias intrinsic to our processing procedure. We also removed clustered SNPs (more than 3 SNPs within a 100bp window).

### Droplet digital PCR

The droplet digital PCR (ddPCR) assays were performed by a QX100™ Droplet Digital PCR System (Biorad) following the manufacturer’s instructions. Each cfDNA sample was mixed with primers and fluorophore labeled probes (FAM for mutant, HEX for wild type). CSF cell pellet DNA was sheared into 300 base pair fragments by sonication (Covaris) before preparing the PCR reaction mix. After PCR, the droplet reader detected fluorescent signals in each droplet.

### LMD and primary tumor mutation retrospective chart review

Charts of a cohort of 212 patients with LMD treated at Stanford Hospital between 2007 and 2017 were examined (see Fig. S1 for cohort selection). We identified 49 NSCLC patients with confirmed LMD. Mutation profiles of these patients’ NSCLC were collected from records of molecular tests performed at Stanford Hospital or another facility. Five patients had no or missing records of molecular panels and were excluded. Analysis to examine the effect of mutation profile on clinical prognosis was carried out on the remaining 44 patients and the 8 WES patients. Of the 44 retrospective patients (29 F, 40-85 yo), 12 were alive at the time of this analysis (Table 1).

**Table 1.** Demographic table of retrospective analysis of 52 LMD patients

|  | **Total**<br>**(n = 52)** | **WES + Retrospective**<br>**EGFR+**<br>**(n = 36)** | **WES + Retrospective**<br>**EGFR-**<br>**(N = 16)** |
| --- | --- | --- | --- |
| **Sex, Female** | 34 (65.4%) | 26 (72.2%) | 8 (50.0%) |
| **age at LMD Diagnosis (year)** |  |  |  |
| Median (Range) | 65 (25 - 85) | 65 (25 - 83) | 64 (40 - 85) |
| **Race/Ethnicity** |  |  |  |
| White | 25 (48.1%) | 16 (44.4%) | 9 (56.3%) |
| Asian | 21 (40.4%) | 16 (44.4%) | 5 (31.3%) |
| Hispanic/Latino | 3 (5.8%) | 3 (8.3%) | 0 (0.0%) |
| Other/Omitted | 3 (5.8%) | 1 (2.8%) | 2 (12.5%) |
| **Smoking History** | 17 (32.7%) | 10 (27.8%) | 7 (43.8%) |
| **Alive at Time of Analysis** | 13 (25.0%) | 7 (19.4%) | 6 (37.5%) |
| **KPS at LMD Diagnosis** |  |  |  |
| Median (Range) | 70 (20 – 90) | 70 (30 – 90) | 65 (20 – 90) |
| **Extracranial Metastases** | 34 (66.7%) | 23 (65.7%) | 11 (68.8%) |
| **Treatment Prior to LMD Diagnosis, %** |  |  |  |
| Chemotherapy | 50 (97.1%) | 34 (97.1%) | 16 (100%) |
| Immunotherapy | 22 (43.1%) | 13 (37.1%) | 9 (56.3%) |
| EGFR Targeted Therapy | 37 (71.1%) | 37 (91.4%) | 0 (0.0%) |
| Brain Radiation | 28 (53.8%) | 17 (47.2%) | 11 (68.8%) |

## Results

### NSCLC LMD mutation characteristics

To identify recurrent genes in LMD without bias, we performed whole-exome sequencing on eight LMD cases to an average sequencing depth of 86X (Fig. S2). Previous studies have shown that lung cancer in never-smokers has a distinct profile of oncogenic mutation different from smokers and former smokers.(19) Six patients in this cohort are never-smokers (Table S1). Normal DNA in saliva or blood samples were obtained from five of the eight patients. The other three patients did not have matched normal tissue. Although the lack of normal tissue complicates somatic variant calling, we reasoned that inclusion of these samples would increase the power to detect recurrent mutation especially given the small cohort size. We used a previously published pipeline to remove germline mutations found in existing public genotype data in 1000 genomes,(17) dbSNP,(20) and the NHLBI Exome Sequencing Project (Exome Variant Server, NHLBI GO Exome Sequencing Project (ESP), Seattle, WA (URL: http://evs.gs.washington.edu/EVS/))(16). We further removed germline variants from ExAC(21) to reduce false positive but retained known oncogenic variants in COSMIC and ICGC to retain likely true positives. To reduce alignment error intrinsic to our pipeline, we removed variants shared between the three unmatched tumor samples and five available normal controls. We compared the genotypes across all sample pairs and detected no sample swap or apparent cross contamination (Fig. S2). All samples harbored between 1034 to 1791 mutations except for LM6, which had 418 mutations (Fig. S3). However, we did not detect any anomaly in distribution of mutation categories for LM6. Amongst all mutations, single nucleotide polymorphisms accounted for 82-94% of each sample, with the rest being small insertions and deletions (Fig. S4). Missense mutations constituted the majority of variants (54.2%), followed by synonymous mutations (34.7%). Mutation profiles were similar across eight samples (Fig. S5). Although synonymous mutations can affect mRNA structure and influence the rate of translation,(22) we only focused on non-synonymous coding variants for ease of interpretation.

### EGFR and TP53 are recurrently mutated in NSCLC LMD

Mutations in *EGFR* were found in six of eight patients (75%). An independent study estimates the *EGFR* prevalence among LMD patients to be 73.8%,(23) in agreement with our study. Deletions between residues 746 and 750 (a.a. sequence Glu-Leu-Arg-Glu-Ala) were found in two patients (LM4, LM8). A similar deletion between residues 747 and 752 (a.a. sequence Leu-Arg-Glu-Ala-Thr-Ser) was found in LM6. A sensitizing recurrent mutation L858R was found in two patients LM1 and LM5. Notably, the somatic mutation in LM5 was initially misidentified by WES due to low mutant allele fraction (Fig. 2A, red dot). We carried out an orthogonal validation with ddPCR on cellular DNA and cell-free DNA from CSF, both of which showed evidence for L858R mutation. A glycine insertion between D770 and N771 on exon 20 was found in LM2. Notably, all mutations were within the tyrosine kinase domain that regulates protein activation (Fig. 2B).(24) We then compared the LMD *EGFR* mutation to clinical mutation panel analyses done on the patients’ primary tumors available for seven out of eight patients. All mutations found in the LMD samples were represented in the primary lung tumors, indicating all metastatic mutations were inherited from the primary tumors.

**Figure 2.**
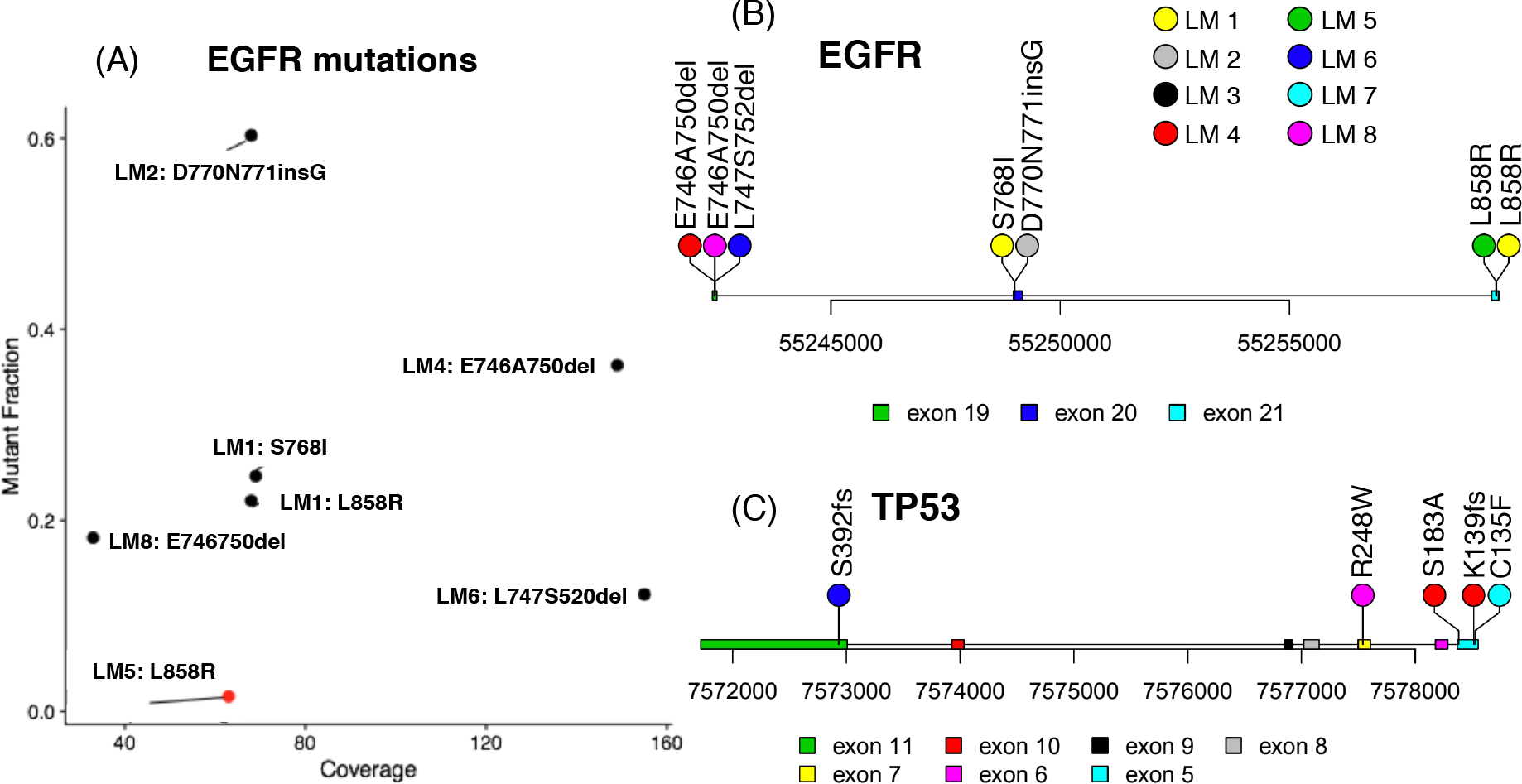
Common gene mutations in LMD identified by WES. (A) Mutation fraction and coverage of each EGFR mutation, LM5 was identified by ddPCR, shown in red; (B) mutation loci on EGFR gene; (C) mutation loci on TP53 gene.

The tumor suppressor gene *TP53* was mutated in four of eight (50%) LMD patients. Of these mutations, one missense mutation S183A and a frameshift mutation K139fs were found in LM4. A frameshift mutation S392fs was found in LM6. A missense mutation C135F was found in LM7. A hotspot missense mutation R248W was found in LM8. Four out of five mutations are located in the DNA binding domain spanning residues 102-292, and the other one (p.S392fs) was located in a regulatory domain spanning 364-393 (Fig. 2C).(25) Among all the *TP53* mutations found in LMD exome seq, only R248W was also in somatic mutation panel test for primary tumor, and it’s not a *de novo* mutation for LM8. We are not able to determine whether any other patient had *de novo TP53* mutation during metastasis due to the lack of overlapping *TP53* mutations in primary tumor mutation panel test and WES.

### Additional NSCLC LMD mutations

A missense mutation E545K in the proto-oncogene *PIK3CA* was found in LM8. The mutation was located within a highly conserved helical domain, and has been shown to promote the catalytic activity of *PIK3CA* resulting in enhanced downstream signaling.(26) A missense mutation S140L in the proto-oncogene *FGFR1* was found in LM7. Although this ultra-rare mutation has a frequency of 0.001318 in the gnomAD database,(21) it was categorized as likely benign by ClinVar.(27) A missense mutation H1180L in the receptor tyrosine kinase *MET* was found in LM5. Although we did not find any previous study on this mutation, it was highly conserved and predicted to be damaging by both SIFT and PolyPhen-2 (Fig. S6). Our data captures mutations overlooked by somatic cancer mutation panels. For instance, WES identified a S289I in PDE4DIP for LM3. This gene is not included in most clinial cancer panels. A comprehensive list of all recurrent mutations that were called in LMD and solid brain metastasis patients is available in the supplementary tabular data S1 and S2.

### KRAS and EGFR mutation frequency differs between LMD and solid metastases

Prior to this study, differences in mutation frequencies between NSCLC solid and LMD brain metastases had not been reported. We processed 26 publically available NSCLC-derived solid brain metastases samples(11) using the same pipeline as our own LMD cohort. We compared the frequency of known recurrent mutations in order to identify shared and unique mutation profiles across LMD and solid brain tumors. Mutations in *TP53* were found in 50% of samples, making it a shared recurrent gene in both types of metastases. *EGFR* showed strong preference towards LMD cases (75% in LMD versus 3.8% in solid brain metastases). To confirm this distinction, we performed a retrospective chart review identifying 44 additional patients with NSCLC and LMD. Together with the eight WES patients, a total of 33 patients (63.5%) were positive for EGFR mutations, much higher than the mutation rate in solid brain tumors (Fisher’s exact test p = 1.518e-07). A mutation in the proto-oncogene *KRAS* was identified in 50% of solid brain metastases but not in any of the LMD cases (Fig. 3). To check if sufficient coverage has been achieved over the *KRAS* region in the LMD samples, we plotted read depth over coding region for LMD cases. The median coverage for coding sequences was between 40 to 134 reads (Fig. S7), which was sufficient to call point mutations. Again, we verified our WES results, through our retrospective case series, a remarkably low mutation rate in lung cancer patients with LMD (4/52 or 7.7%), much lower than the mutation rate in solid brain metastasis (Fisher’s exact test, p = 4.85e-05). Likewise, given the expected *KRAS* mutation frequency is around 30%^25^ in lung cancer, the lower frequency of *KRAS* mutations in LMD patients is notable.

**Figure 3.**
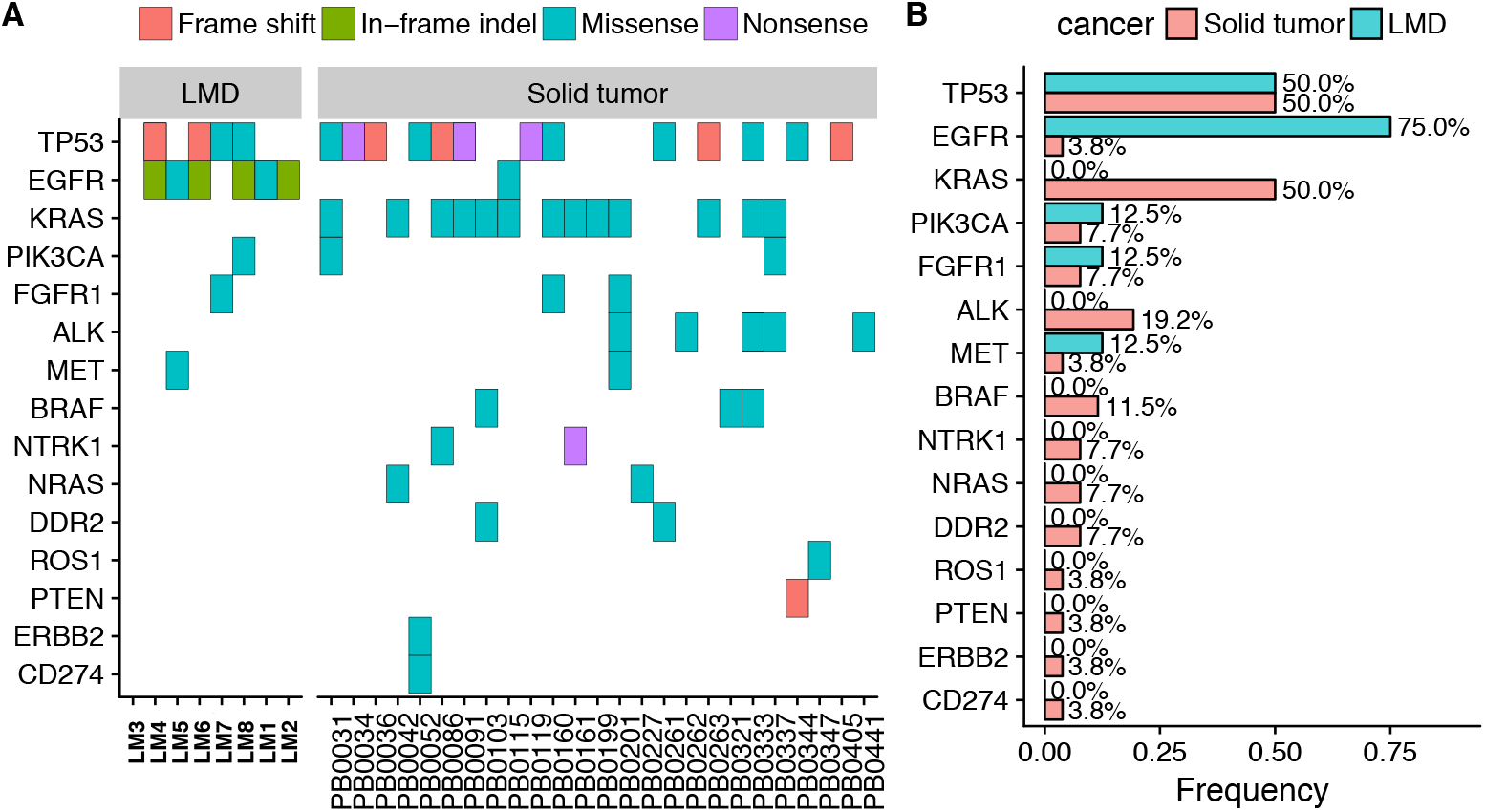
Canonical genes in LMD and solid brain metastases. (A) mutation types across all samples (B) mutation frequency in LMD and solid metastases

### NSCLC LMD cell-free DNA detected in CSF

Brain tumor-associated cfDNA has been previously detected in CSF.(28,29) Primary lung cancer routinely undergoes clinical mutation testing to determine the most appropriate targeted therapeutic, as shown in Table S1. Droplet digital PCR assays were performed to validate the point mutations found in WES and investigate NSCLC LMD cfDNA in CSF. Here we show two canonical mutations found in the primary lung tumor, which we validated in CSF via ddPCR. The primary lung tumor of LM5 was known to have *EGFR^L858R^* mutation, and the mutation was detected in LM5 cfDNA (Fig. 4A) and cellular DNA (Fig. 4B) from the patient’s CSF. Droplet digital PCR revealed a mutant allele fraction of 6.4% in cellular DNA and 25% in cfDNA. Meanwhile, in the exome sequencing of cellular DNA, only one out 63 reads showed a T>G mutation, falling below our pipeline’s calling threshold. The primary lung cancer of LM8 was known to have a point mutation *TP53^R248W^*, and the mutation was validated in CSF cfDNA (Fig. 4C) and cellular DNA (Fig. 4D). In this sample, the mutant allele fraction in cellular DNA was high (95%), which is consistent with exome sequencing (10 out of 11 reads showed mutation).

**Figure 4.**
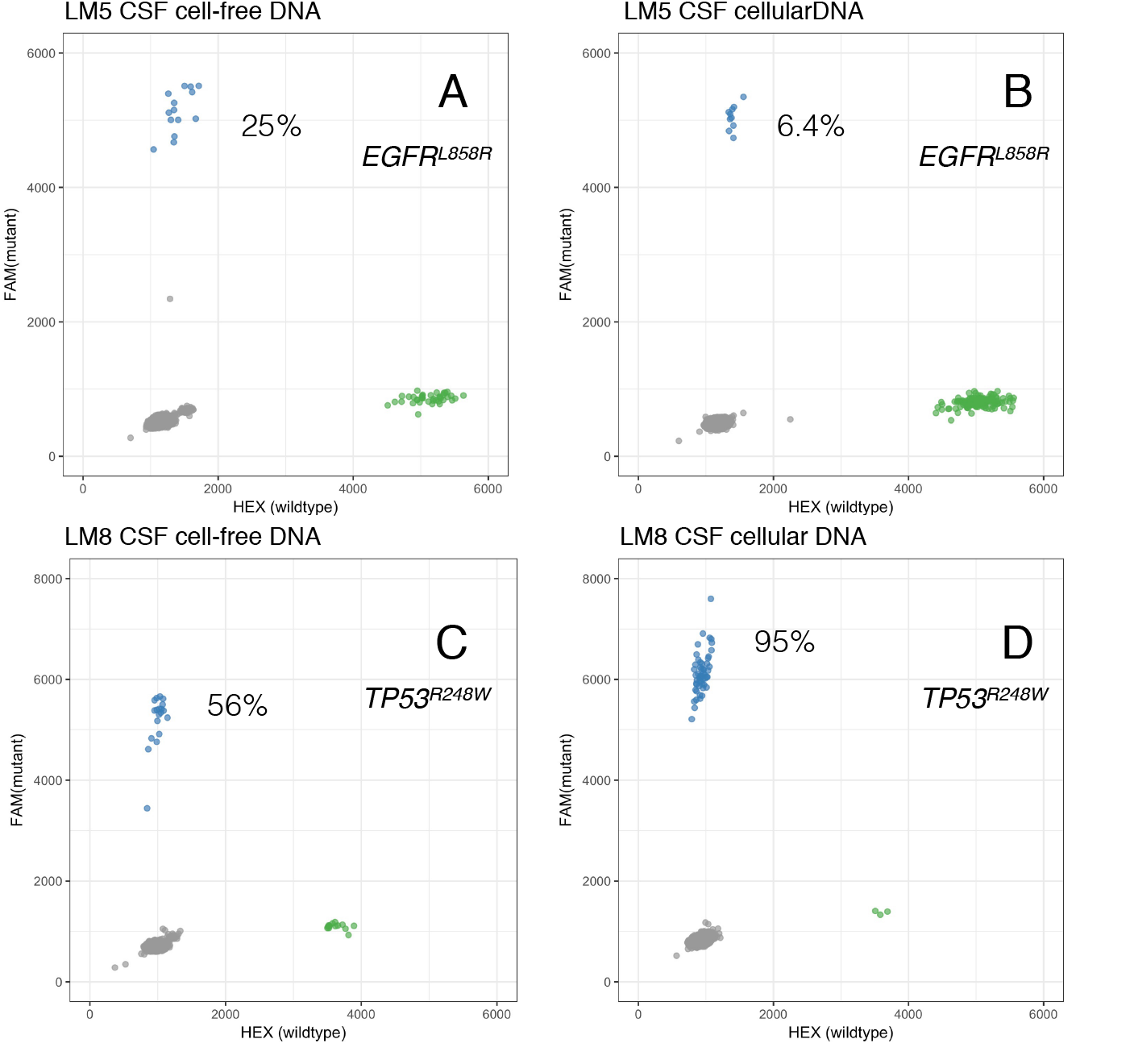
Cell free DNA in cerebral spinal fluid reflects mutations in NSCLC leptomeningeal disease. Representative ddPCR plots of point mutations detected in CSF cellular DNA and cfDNA in two patients with cytology positive CSF. LM5 (A, B) underwent ddPCR for *EGFR*^L858R^, while LM8 was tested for mutation *TP53*^R248W^. The mutant fraction in cell free (A, C) compared to cellular (B, D) DNA varies depending upon peripheral blood contamination.

### Higher prevalence of EGFR mutation in LMD cohort

The genetic data suggested that *EGFR* mutations were more common in our LMD cohort. We investigated the correlation of EGFR mutations with LMD vs solid lung-to-brain metastases. Of our 44 retrospectively reviewed patients (Fig. S1) and our 8 WES patients, 33 LMD patients had a positive EGFR mutation. We then performed a Wilcoxon rank sum test to compare the time between primary lung tumor and LMD diagnosis. Patients with *EGFR* mutations had a median interval of 16.3 months, compared with 23.9 months for patients without EGFR mutations (Fig 5A; p = 0.083). Kaplan-Meier curves on interval between primary lung and LMD diagnoses suggested patients with EGFR mutation showed a faster progression towards LMD, although this result was not significant (Fig 5B; p = 0.0863). This analysis is likely underpowered given the small numbers of patients in our cohort. It is known that patients with EGFR mutations benefit from tyrosine-kinase inhibitor (TKI) treatment as first line therapy.(23) To correct for these potential confounding effects of differences in treatment, we used a Cox proportional hazard model with sex, race, age at primary diagnose, KPS score, smoking history and treatment (immunotherapy, brain radiation, targeted therapy, and tyrosine-kinase inhibitors) as covariates. *EGFR* mutation significantly increased the hazard (p = 0.017), whereas targeted therapy significantly decreased the hazard (p = 0.035).

**Figure 5.**
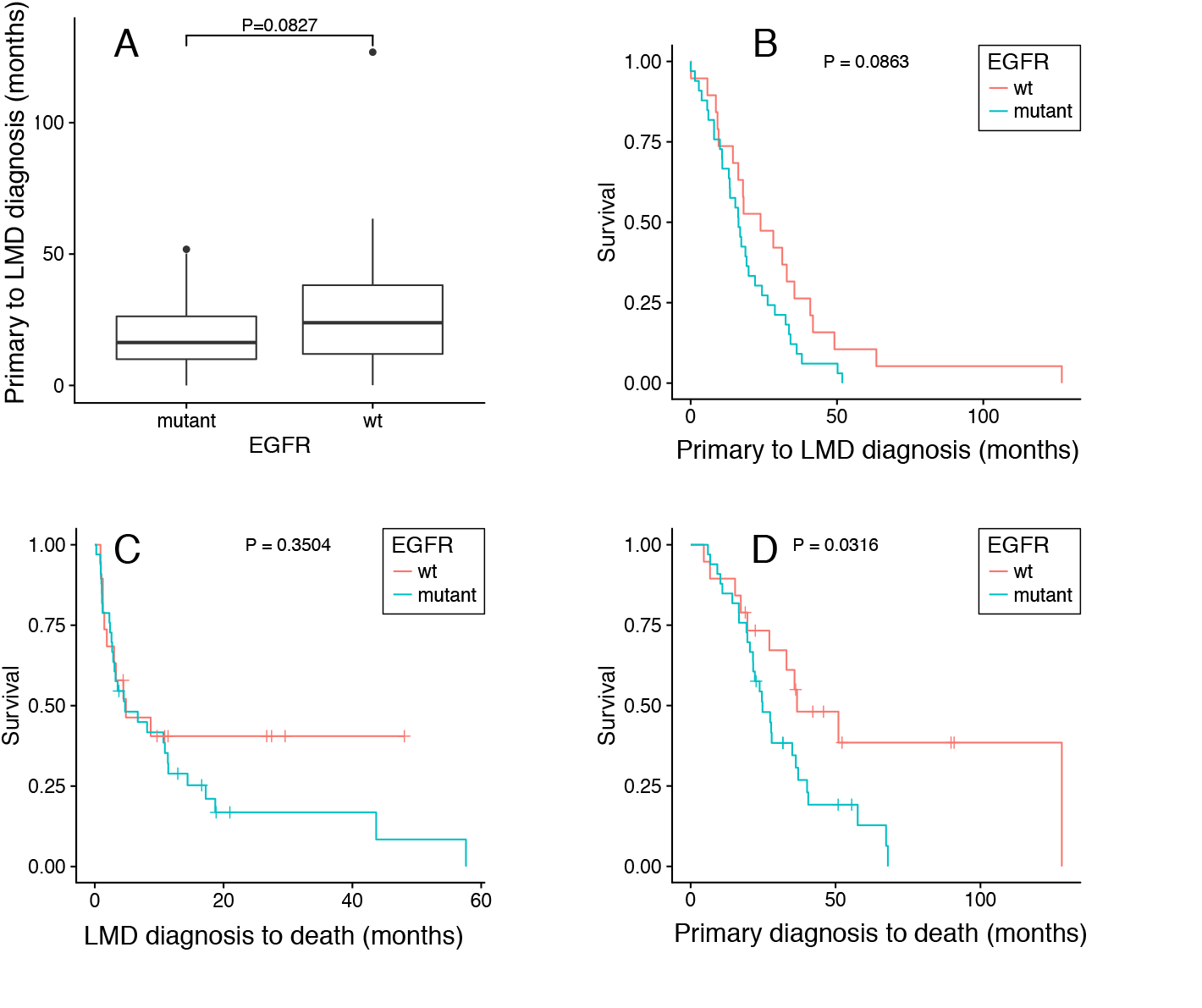
Correlation of *EGFR* mutation and LMD progression. (A) Wilcoxon rank sum test of time interval from primary cancer diagnosis to LMD diagnosis (B) Kaplan-Meier plot on interval from primary cancer diagnosis to LMD diagnosis. (C) Patients survival from the point of LMD diagnosis; (D) Patients survival from the point of primary cancer diagnosis.

## Discussion

NSCLC leptomeningeal and solid brain metastases have distinct mutation profiles

Overall, the most common recurrently mutated genes in our study were *TP53, EGFR*, and *KRAS*. Interestingly, *EGFR* mutations occurred more frequently in the LMD samples while *KRAS* mutations occurred more frequently in solid brain metastasis samples. This finding was supported by a retrospective chart review with an additional set of 44 LMD patients, only 4 (7.7%) of whom had *KRAS* mutation. In contrast, 33 (63.5%) patients had *EGFR* mutations. An earlier retrospective study from our institution found that *KRAS* mutations were present in 2 of 30 (7%) and *EGFR* mutations were present in 13 of 30 (43%) of NSCLC LMD patients.(7) *EGFR* and *KRAS* are mutated in approximately 17%(30) and 30%(31) of patients with lung adenocarcinoma, respectively. *EGFR* mutations and *KRAS* mutations are believed to be mutually exclusive, though concomitant mutations can occur rarely.(32)

The prominence of *EGFR* mutations and lack of *KRAS* mutations in LMD suggests intrinsic differences in tumor biology that allow these metastatic cells to thrive in distinct brain niches, or selection by progressive systemic treatments for subpopulations of cells from the primary tumors that are particular to a brain niche. One hypothesis we considered was that patients with *EGFR* mutations treated with TKIs survived longer(33) and thus had more time to develop LMD, whereas patients with *KRAS* mutations were not responsive to TKIs and typically had worse prognosis.(34) However, data from our LMD cohort did not support this hypothesis, as patients with *EGFR* mutations either similar survival time from the point of LMD diagnosis (Fig. 5C) or shorter survival time from primary cancer diagnosis to death (Fig. 5D). A Cox model likewise did not find *EGFR* mutations to be associated with survival time after LMD diagnoses (p = 0.16). *TP53* was mutated in 50% of both LMD and solid lung cancer brain metastasis samples, suggesting that disruption in *TP53* does not prefer either brain niche. In a recent study by Villaflor *et al* of 68 patients with NSCLC, circulating tumor DNA was sequenced to identify actionable tumor mutations in plasma.(35) The most commonly mutated genes was *TP53* (40% of patients),which is largely consistent with our result. Mutations in *PIK3CA, ALK, MET, FGFR1*, and *BRAF* are known to occur in lung cancer at much lower rates,(35) and this was reflected as well in our patients with solid NSCLC brain metastases.

The paucity of data on LMD and its rare occurrence make recurrent mutation discovery challenging and difficult to interpret. Our analysis also found several other genes not listed as recurrent gene in the current literature. For instance, *TAS2R31*, a bitter taste receptor gene, is mutated in 4 of 8 LMD samples (50%) and in 1 of 26 (3.8%) solid lung cancer samples. The significance of these mutations is difficult to ascertain as they are not well described in the literature. One recently published work compared recurrently mutated genes in primary lung adenocarcinoma and matched chest wall metastasis nodules in four patients.(36) Interestingly, *TAS2R31* was mutated in 2 of 4 of the metastatic tumors and in none of the primary tumors.

### Droplet Digital PCR preferable to WES in detecting LMD-associated DNA in CSF

WES could provide a comprehensive genetic analysis of an LMD patient sample. However, due to the exome-wide coverage, the detection limit is often compromised when low levels of DNA are present. This is especially concerning with our cohort given the amount of DNA across LMD CSF samples is highly variable. For example, cytology positive CSF could have cell numbers ranging from several to tens of thousands. In our study, the DNA extracted from the pellets ranged from 20 ng to 200 ng, and cfDNA levels ranged from 0.5 to 5 ng. In addition, CSF is generally obtained through lumbar puncture or ommaya reservoir, frequently with some peripheral blood contamination. This can vary unpredictably the tumor DNA fraction from the cell pellets and poses a problem in exome sequencing for variant detection.

We have previously found ddPCR to be the most sensitive method for tumor-associated DNA detection in CSF(37). The source of cfDNA includes cell apoptosis, tumor cell necrosis and secretion.(38) As there should be no cells in the CSF of healthy individuals, the background noise of normal DNA is low, making cfDNA a good diagnostic tool with higher sensitivity than regular cytology. We found that low DNA levels present in LMD samples favor ddPCR over WES for variant detection. For example, in LM5, ddPCR showed 6.4% mutant allele fraction of *EGFR^L858R^* in cellular DNA, a lower mutant allele fraction than the cell-free component (25%), a difference likely due to blood contamination. In comparison, WES analysis did not identify the *EGFR* mutation, likely representing a false negative. On the other hand, in sample LM8, the mutant allele fraction of *TP53^R248W^* mutation in cellular DNA calculated from ddPCR (95%) was consistent with WES data (91%). In contrast to LM5, this patient had extremely high tumor cell count in CSF at the time of sampling (~10^5^ cells total), which is rare. In this case, the cell-free component had lower mutant allele fraction (56%) than cellular DNA. This contrast would suggest caution when generating conclusions from WES data with low amounts of input DNA, given the lower coverage and risk of false negative results.

### EGFR and LMD Progression

*EGFR* mutation frequency in our LMD cohort (63.5%) was higher than that in lung cancer patients overall (15%~30%).(30) Similar results were seen by Li *et al*. with an *EGFR* mutation frequency of 73.8%(23) in their cohort of 160 NSCLC patients with LMD. A retrospective review of 1,127 NSCLC patients(39) showed EGFR-mutated cases to have a higher prevalence of brain metastases (31.4% vs 19.7%) and likelihood of leptomeningeal dissemination (30.8% vs 12.7%). Our data suggest a correlation between *EGFR* mutations and disease progression from primary lung cancer to LMD (Fig. 5B). Patients with EGFR-mutated NSCLC developed LMD faster than wild-type patients. *EGFR* mutations have been previously shown to occur early in NSCLC pathogenesis and to increase in frequency through cancer progression to advanced metastasis.(40–42) In a cohort of NSCLC patients with solid brain metastases, Matsumoto and colleagues found retention of *EGFR* mutation profile between the primary and metastatic tumors, consistent with our LMD cohort.(42)

While the benefit of targeted therapeutics on overall survival is well established, its effect on the development of LMD and the prognostic relationship once LMD has been diagnosed is still under investigation(43,44). In our study, targeted therapeutics slowed the progression of primary lung disease to LMD. However, regardless of targeted treatment, *EGFR* mutation accelerated progression to LMD. Our data suggest selective treatment pressure, and not overall survival, accounts for the high percentage of patients in our LMD cohort with *EGFR* mutations. One consideration is the relative lack of permeability to the blood-brain and blood-tumor barrier which may allow for a haven for tumor cells in the leptomeninges. EGFR-mutated NSCLCs may metastasize within this protective space prior to treatment. In contrast, LMD from non-EGFR-mutated NSCLCs such as KRAS (7.7% of our cohort vs the expected 30%^25^) was less often observed in LMD.

## Conclusion

In contrast to solid brain metastases that have excellent local control rates following surgery and radiation, LMD remains nearly universally and rapidly fatal. The development of effective LMD therapies requires a greater understanding of LMD genomics and cancer biology. NSCLC solid and LMD brain metastases have distinct mutation profiles which may correlate with either an intrinsic propensity to grown within these distinct brain niches, or selective treatment pressures placed through exposure to different systemic therapies. The overrepresentation of *EGFR* mutations and relative lack of *KRAS* mutations in our LMD cohort reflect this question of genetic predisposition to metastasize to the leptomeninges and thrive despite systemic treatment exposure. Our data suggest that mutations in *EGFR* and *TP53* hold great potential as biomarkers for LMD diagnostics, as a mutation in one or both genes was detected in every patient in our cohort. Given the rarity of LMD, we are committed to facilitate sharing our sequencing data with the greater scientific community, and we have deposited these results in a readily accessible website at www.LMDseq.org.

## Acknowledgement

We acknowledge the generous donation of clinical samples by our patients, without which this research would not be possible, and to whom we dedicate our work. Funding to MHG included R21CA193046-01, K08NS901527, the Hearst and Curci Foundations. YL is supported in part by the McCormick-Gabilan Award and Stanford CHRI fellowship; BL is supported by the Stanford CEHG fellowship and the National Key R&D Program of China, 2016YFD0400800. IDC is supported by the Stanford Medscholars Grant. We thank Norma Neff and Gary Mantalas for sequencing support, and Tej Azad for providing two clinical samples.

## Supplementary Material

Supplementary material is available online.

